# PARP1 Writes N3-Cytidine ADP-Ribosylation in DNA

**DOI:** 10.64898/2025.12.18.695099

**Authors:** Michael U. Musheev, Lars Schomacher, Johanna M. Schott, Amitava Basu, Martin M. Möckel, Sabine Heinen, Laura Frosch, Peng Guo, Gaofan Yang, Qiuya Huang, Christof Niehrs

**Affiliations:** Institute of Molecular Biology (IMB), 55128 Mainz, Germany; Chengdu ChemPartner, Chengdu 610041, China; DKFZ-ZMBH Alliance, Division of Molecular Embryology, 69120 Heidelberg, Germany

**Keywords:** PARylation, DNA modification, Mass spectrometry, Enzyme catalysis, Nucleic acids

## Abstract

Recent evidence indicates that mono - and poly-ADP ribosylation (MARylation and PARylation) are not limited to proteins but extend to DNA. Notably, *in vitro* base PARylation by PARP1 in single stranded DNA (ssDNA) was demonstrated at N1-deoxyadenosine (N1-dA). Here, we report that PARP1 catalyzes N3-specific ADP-ribosylation of deoxycytidine (N3-dC) in single-stranded DNA. Analogous to N1-dA PARylation, which is prone to spontaneous adenine-to-inosine deamination, N3-dC PARylation promotes cytosine deamination, yielding N3-PARylated-deoxyuridine. These deamination products yield diagnostic PARylation signatures in LC–MS/MS, namely N1-ribosyl-deoxyinosine (N1-r-dI) and N3-ribosyl-deoxyuridine (N3-r-dU). We synthesized both N1-r-dI and N3-r-dU as diagnostic standards and established absolute quantification of base ADP-ribosylations by LC–MS/MS. Quantitative analysis of PARylated dA and dC in ssDNA reveals pronounced sequence preferences of PARP1. Removal of these base modifications differs markedly, since ADP-ribose glycohydrolase TARG1 removes PAR from both dA and dC, whereas PARG acts exclusively on dA. Our results establish cytidine ADP-ribosylation as a novel DNA modification, with potential roles in DNA metabolism, epigenetic regulation, or genome stability.

## Introduction

ADP-ribosyltransferases (ARTs) are well known to post-translationally modify proteins by mono- and poly-ADP ribosylation (MAR- or PARylation), but only recently were they shown to modify also RNA and DNA in a variety of organisms^1^. In butterflies, mollusks and Streptomyces, the enzyme-toxins pierisin, CARP-1 and Scabin MARylate the N2 position of guanosine residues in DNA^2–4^. In butterflies, DNA-MARylation may potentially serve as a defense factor against parasitism^5^. In bacteria, DarT1 and DarT2 are part of toxin-antitoxin systems that in ssDNA MARylates guanosine at N2 and thymidine at N3, respectively^6,7^. In mammals, ARTs modify *in vitro* phosphorylated 3’- and 5’-termini of DNA by PARylation (PARP1 and PARP2) and MARylation (PARP3)^8–11^, and *in vivo* at telomeres^12^. Recently, we reported that PARP1 modifies single stranded DNA (ssDNA) *in vitro* at the N1-position of adenosine and showed that the modification naturally occurs in mammalian DNA^13^. These findings underscore the need to establish a synthetic route to obtain chemical standards for absolute quantification of N1-dA ADP-ribosylation in genomic DNA. Moreover, the findings raise the possibility that base- ADP-ribosylation may not be limited to N1-dA.

Here we address both issues, showing PARP1 modifies ssDNA *in vitro* also at cytosine-N3 and reporting the synthesis of chemical standards that enable absolute quantification of both dA- and dC- ADP-ribosylation by LC-MS/MS. We use these chemical standards to confirm the natural occurrence of DNA base ADP ribosylation in mammalian cells and to quantify its levels.

## Results

Since PARP1 modifies ssDNA *in vitro* at N1-dA^13^, we asked if other canonical DNA bases could also function as ADP-ribose acceptors and focused on the N3 of dC, which, like N1-dA, lacks a proton in its standard tautomeric form. Specifically, we screened by LC-MS/MS for the occurrence of ribosyl-deoxyuridine (N3-r-dU) as diagnostic molecule with the following reasoning. PARylated N1-dA is prone to deamination when degraded to single nucleosides during LC-MS/MS work up. Hence, the diagnostic nucleoside for dA-PARylation is its ribosylated deamination product, ribosyl-deoxyinosine (N1-r-dI)^13^. Similarly, alkylation of N3-dC destabilizes the aromatic ring, and favours spontaneous deamination at physiological pH^14^. Hence, potential N3-dC PARylation may also promote deamination of PARylated dC into PARylated dU (**Fig. 1a**). Upon DNA cleavage, the diagnostic molecule to monitor dC-PARylation by LC-MS/MS would be N3-r-dU.

**Figure 1:**
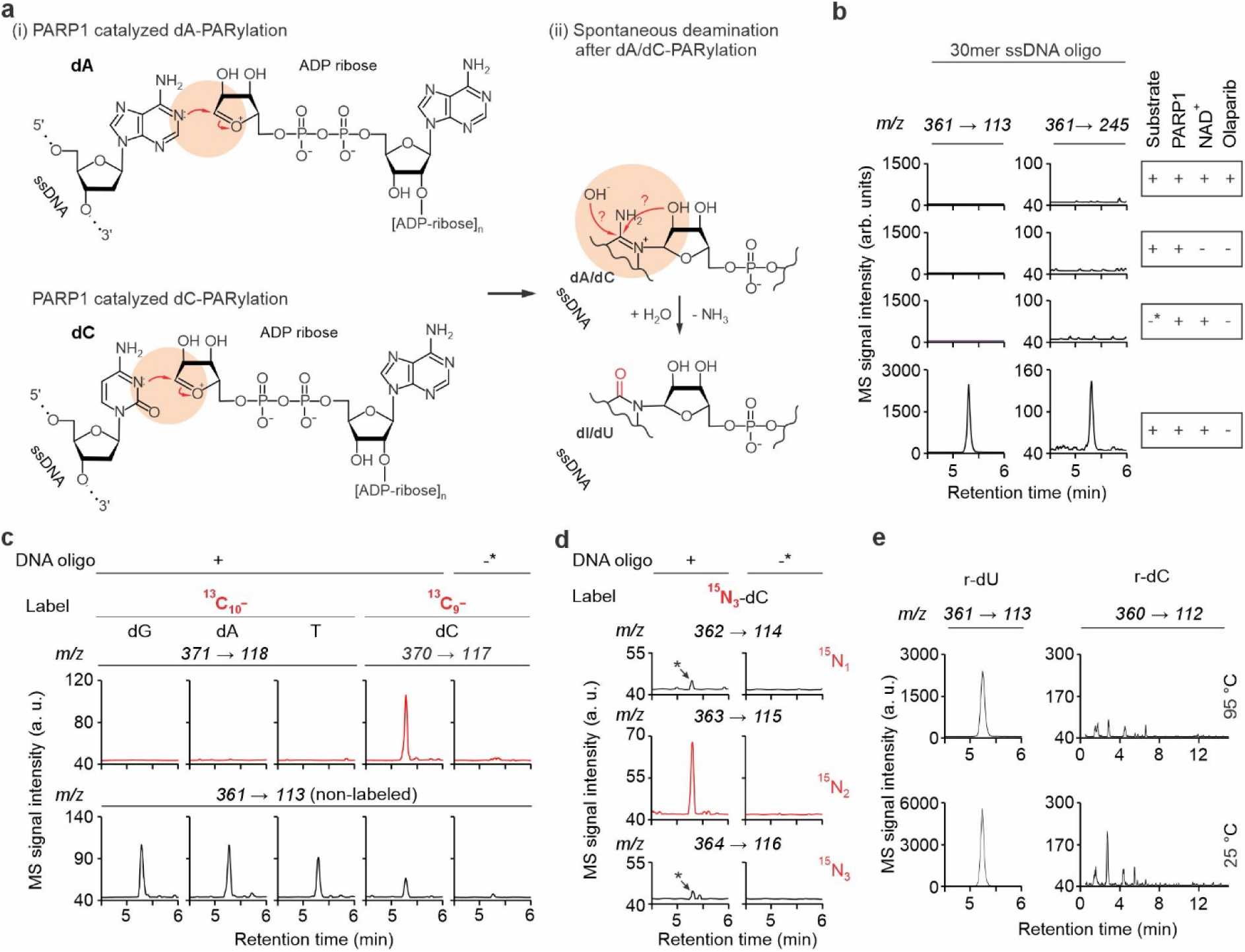
ADP ribosylation of cytidine in ssDNA by PARP1. (a), Proposed mechanism of (i) N1-adenosine and N3-cytidine ADP-ribosylation in ssDNA by nucleophilic attack of the ring-nitrogen towards the electrophilic C1’-carbon of the ribose, and (ii) spontaneous deamination of PARylated N1-dA and N3-dC. Note that the chemical reactions are identical for dA and dC. (b), LC-MS/MS chromatograms of reaction products from *in vitro* PARylation with PARP1 in presence or absence of NAD^+^, the PARP inhibitor Olaparib, and a 30mer ssDNA standard oligonucleotide as indicated. Left and right panels show mass transitions corresponding to mass shifts expected for loss of a deoxyribose + ribose (m/z 361 → 113) and loss of a single deoxyribose (m/z 361→ 245) from the parental ‘ribosyl-dU’ (r-dU); *, 30mer oligonucleotide was added after phenol inactivation of PARP1. (c), LC-MS/MS chromatograms as in (b) but from *in vitro* PARylation with PARP1 on four 83mer ssDNA oligos each labeled with one type of heavy carbon containing nucleoside as indicated. Top, scan for signals with m/z transitions expected for a mass shift of r-dU from each label. Bottom, detection of signals that correspond to non-labeled r-dU. (d), LC-MS/MS chromatograms as in (b) but from *in vitro* PARylation with PARP1 on ^15^N_3_-dC-labeled 83mer ssDNA oligo. Reaction products were scanned for signals with m/z transitions that correspond to mass shifts of +1 to +3 Da of r-dU (from top to bottom). Arrows mark signals of natural occurring isotopologues of r-dU and ^15^N_3_-r-dU. Note that (i) a 83mer oligonucleotide length, rather than the 30mer used for the standard oligonucleotide in (b), was used for heavy isotope labeling by PCR, and (ii) the labeled 83mer oligonucleotides in (c) and (d) contained a mixture of labeled and unlabeled nucleosides, allowing detection of both labeled and unlabeled reaction products. (e), LC-MS/MS chromatograms from *in vitro* PARylation assays as in (b) in which the reaction products were either processed at 95 °C or 25 °C before mass spec analysis, and screened for signals with m/z transitions expected for r-dU (m/z 361 → 113, left) and r-dC (m/z 360 → 112, right).

We performed *in vitro* ADP ribosylation of a 30mer ssDNA oligonucleotide (“30mer”) that is PARylated by PARP1^13^, degraded the DNA to nucleosides, and monitored for r-dU by LC-MS/MS. Indeed, we observed a positively charged ion with a m/z of 361, matching r-dU. Moreover, mass transitions that fit to loss of a ribose and a deoxyribose (m/z = 361 → 113), as well as loss of a single deoxyribose (m/z = 361 → 245) supported the presence of both sugars on the nucleobase (**Fig. 1b**). Importantly, the loss of a deoxyribose while ribose remains attached reflects N-glycosidic bond cleavage during MS/MS; this fragmentation retains ribose only when it is base-linked, whereas a sugar-linked ribose would be co-lost with deoxyribose, consistent with genuine base PARylation. Detection of r-dU was dependent on the simultaneous presence of PARP1, the 30mer oligonucleotide, and NAD^+^, and was blocked by the PARP1 inhibitor olaparib. Next, we confirmed that r-dU originated from PARylated dC using ssDNA oligonucleotides that each contained one isotopically labelled nucleoside (^13^C_10_-dG, ^13^C_9_-dC, ^13^C_10_-T or ^13^C_10_-dA). Upon *in vitro* PARylation, only the ^13^C_9_-dC-labeled oligonucleotide led to the detection of a mass-shifted molecule, specifically of 361 + 9 Da, as expected for isotopically labelled r-dU (**Fig. 1c**). Of note, the dA-PARylation diagnostic molecule N1-r-dI and its labeled ^13^C_9_-dA analogue appears in a separate channel (m/z = 395 → 142)^13^.

If PARylation of dC promotes its deamination either within intact DNA or during LC-MS/MS work up of hydrolysis products, a PARylated oligonucleotide substrate labelled with ^15^N_3_-dC should lose one nitrogen, resulting in a 361 + 2 instead of +3 Da shift of the diagnostic molecule, which was confirmed (**Fig. 1d**). Interestingly, we did not detect r-dC, deriving from the presumed primary dC-PARylation product, even in freshly prepared samples and omitting heat denaturing prior to DNA degradation (**Fig. 1e**). In contrast, r-dA was readily detectable in non-heated samples (**Extended Data Fig. 1**). The result suggests rapid deamination of PARylated-dC to occur already within intact DNA. Taken together, these results support N3-dC being the ADP ribose acceptor site.

To unambiguously corroborate this conclusion and to obtain a standard, we set out to synthesize N3-r-dU via N-ribosylation of the nucleobase using a Mitsunobu-type (n-Bu₃P/ADDP) protocol (**Extended Data Fig. 2, Fig. 2a**) and obtained the desired compound. Importantly, LC-MS/MS mass transition as well as retention time of the N3-r-dU standard matched the r-dU molecule generated by *in vitro* PARylation (**Fig. 2b**). Furthermore, a mixture of the synthetic and the enzymatically generated molecules co-eluted in LC-MS/MS, supporting their structural identity (**Fig. 2b**). By analogous route and similar yield, we also obtained a synthetic standard for N1-r-dI (**Extended Data Fig. 2,** and **Fig. 2c**), which co-eluted with the degradation product of *in vitro* PARylated N1-dA residues (**Fig. 2d**).

**Figure 2:**
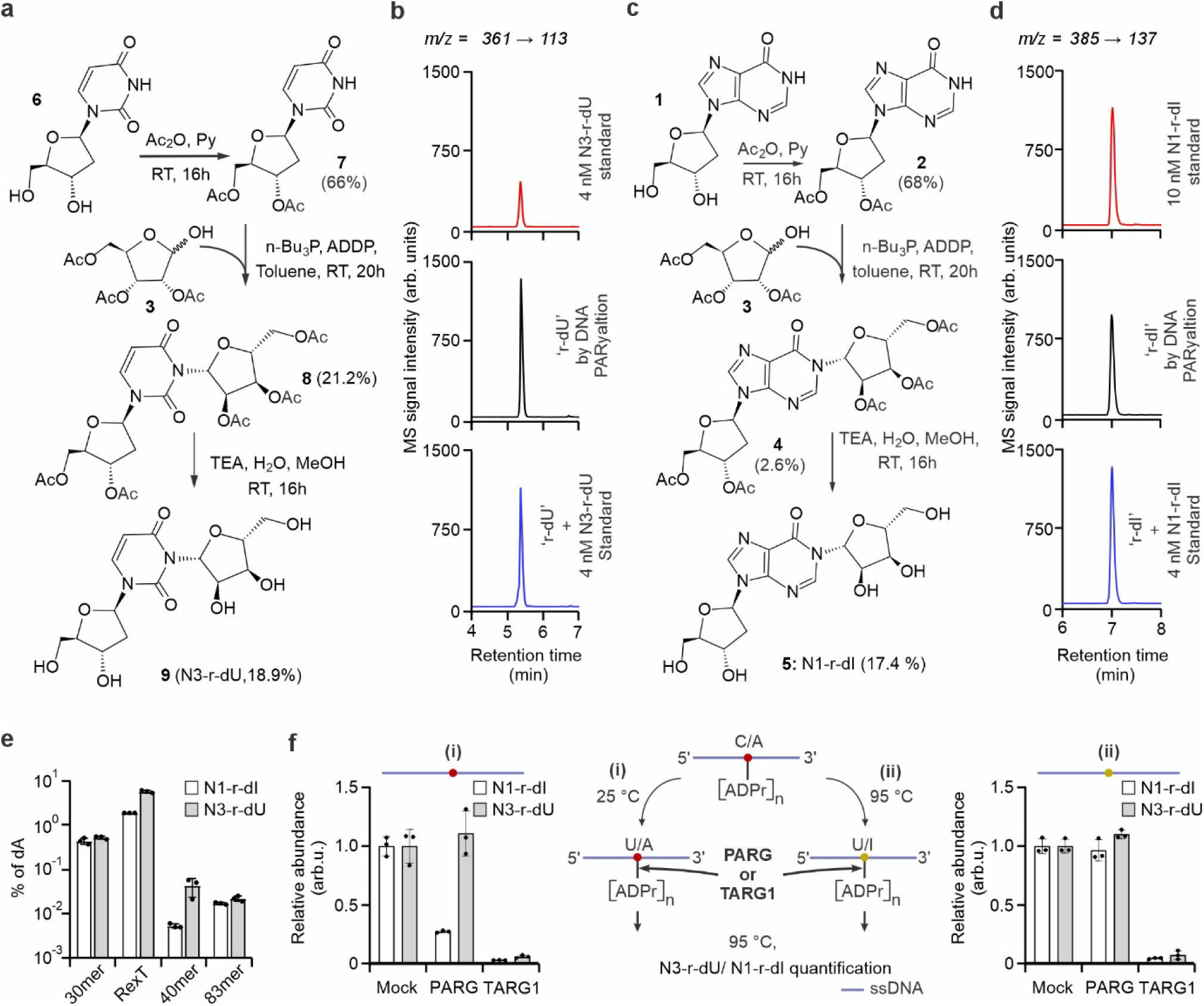
Chemical synthesis of N3-r-dU and N1-r-dI standards and quantification of dA-and dC-PARylation. **a, c**, Synthesis route for N3-r-dU N3-r-dI (see Supporting Information for details). **b**, LC-MS/MS electropherograms corresponding to m/z of N3-r-dU ions (m/z=361→ 113) for synthetic N3-r-dU (top), for r-dU enzymatically generated by PARP1 (middle), and a mixture of equal volumes of the two above mentioned samples (bottom). **d**, LC-MS/MS electropherograms corresponding to m/z of N1-r-dI ions (m/z=385→137) for synthetic N1-r-dI (top), for r-dI enzymatically generated by PARP1 (middle), and a mixture of equal volumes of the two above-mentioned samples (bottom). **e**, LC-MS/MS-based quantification of N1-r-dI and N3-r-dU normalized to dA in different ssDNA oligonucleotides as indicated (s.d., n = 3 independent reactions). **f**, LC-MS/MS-based relative abundance of N1-r-dI and N3-r-dU of an *in vitro*-PARylated 30mer oligo either non-heated (left, condition (i)) or heated at 95 °C (right, condition (ii)) and subsequently treated with PARG and TARG1, respectively (s.d., n = 3 independent reactions).

Next, to enable absolute quantification of both dA- and dC-PARylation in DNA using stable isotope dilution LC-MS/MS, we synthesized the deuterium-substituted isotopologues D_4_-N1-r-dI and D_4_-N3-r-dU using the routes shown in **Extended Data Fig. 3**. Using these standards, stable isotope dilution LC-MS/MS showed a limit of quantification (LOQ) of one and three fmol for N1-r-dI and N3-r-dU, respectively (**Extended Data Fig. 4**). We quantified levels of dA- and dC-PARylation after *in vitro* PARylation of four ssDNA oligonucleotides that differed in length and sequence. Under the conditions used, N1-r-dI and N3-r-dU levels ranged from ∼0.002 – 2 % of total dA (**Fig. 2e**). Notably, the oligonucleotides employed in this study have arbitrary DNA sequence and modification levels may be considerably higher in physiologically relevant sequences. The yields of N1-r-dI and N3-r-dU were proportional to one another across the tested sequences, suggesting that PARP1 does not prefer dA or dC. Together with the broad range of product levels, the result indicates that base PARylation is strongly influenced by DNA secondary structure.

We previously demonstrated that N1-dA PARylation is reversible by the PAR glycohydrolase PARG^13^. Accordingly, LC-MS/MS analysis of *in vitro* PARylated ssDNA showed that PARG treatment reduced levels of N1-r-dI (albeit incompletely), however, N3-r-dU levels were unchanged (**Fig. 2f, condition (i)**), indicating differential substrate specificity towards dA- and dC-PARylation. By contrast, terminal ADP-ribose glycohydrolase TARG1 markedly reduced both N1-r-dI and N3-r-dU (**Fig. 2f, condition (i)**).

What underlies the differential PARG activity towards dA- and dC-PARylation? N1-PARylated dA partially deaminates to N1-PARylated dI in ssDNA (∼20% within 5 days at 4 °C)^13^, whereas *in vitro* PARylation of dC lead to complete deamination to N3-PARylated dU (**Fig. 1e**). In both cases, the N-glycosidic nitrogen (purine N1; pyrimidine N3) is neutral (**Fig. 1a**). This suggests that PARG-mediated hydrolysis requires a protonated nucleobase nitrogen at the N-glycosidic linkage, explaining why the deaminated adducts N1-ADPr-dI and N3-ADPr-dU, which possess a neutral N, are not processed (**Fig. 2f, scenario (i)**). To test this hypothesis, we induced complete deamination of PARylated dA to PARylated dI by heating an *in vitro* PARylated 30mer prior to PARG treatment. Under these conditions, PARG was unable to remove the terminal ADP ribose from N1-PARylated dI (**Fig. 2f, condition (ii)**), confirming its specificity for non-deaminated PARylated bases. In contrast, TARG1 efficiently demodified base PARylation regardless of the base deamination (**Fig. 2f, condition (ii)**).

N1-dA ADP-ribosylation occurs naturally in mammalian DNA, albeit at very low levels^13^. To test for N3-dC ADP-ribosylation *in vivo*, we enriched ssDNA from pig liver total DNA with the diagnostic standards for quantification of dA- and dC- ADP-ribosylation (D4-N1-r-dI and D4-N3-r-dU) spiked-in. We corroborated the occurrence of N1- ADP-ribosylated dA in genomic DNA at levels of ∼6 modifications per genome (**Fig. 3**). This value represents a population average and could be substantially higher if the modification is confined to a small subpopulation of cells. In contrast, LC-MS/MS signals corresponding to N3-r-dU, i.e. N3-dC ADP-ribosylation, were below the detection limit (**Fig. 3**). DNA ADP-ribosylation may be under tight control by PAR glycohydrolases and hence highly transient^12^, possibly explaining low or non-detectable levels of base ADP-ribosylation in primary tissues. Hence, we used gene editing to create mouse embryonic stem cells (mESCs) in which *TARG1* was inactivated and that additionally carry an auxin-inducible PARG degron, for either single TARG1 or simultaneous TARG1/PARG depletion (**Extended Data Fig. 5**). We prepared total DNA from these mESCs, enriched for ssDNA, spiked in D_4_-N1-r-dI and D_4_-N3-r-dU, and monitored for N1-r-dI and N3-r-dU by LC-MS/MS. Interestingly, in the absence of TARG1 but presence of PARG, we could neither detect N1-r-dI nor N3-r-dU while the spiked-in standards were recovered, excluding large losses during sample processing (**Fig. 3**). However, in the absence of both TARG1 and PARG (Auxin-treatment) we detected N1-r-dI at levels that correspond to ∼2 ADP-ribosylation dA-residues per mESC genome (**Fig. 3**). The result demonstrates a PARG-dependent dynamic turnover of dA- ADP-ribosylation in mESCs *in vivo*. In contrast, and consistent with the lack of terminal glycohydrolase activity of PARG toward N3- ADP-ribosylation dU *in vitro*, PARG-depletion in TARG1-deficient mESCs did not yield detectable N3-r-dU. We conclude that unlike N1-dA- ADP-ribosylation, N3-dC- ADP-ribosylation is not readily detectable in pig liver tissue and mESCs.

**Figure 3:**
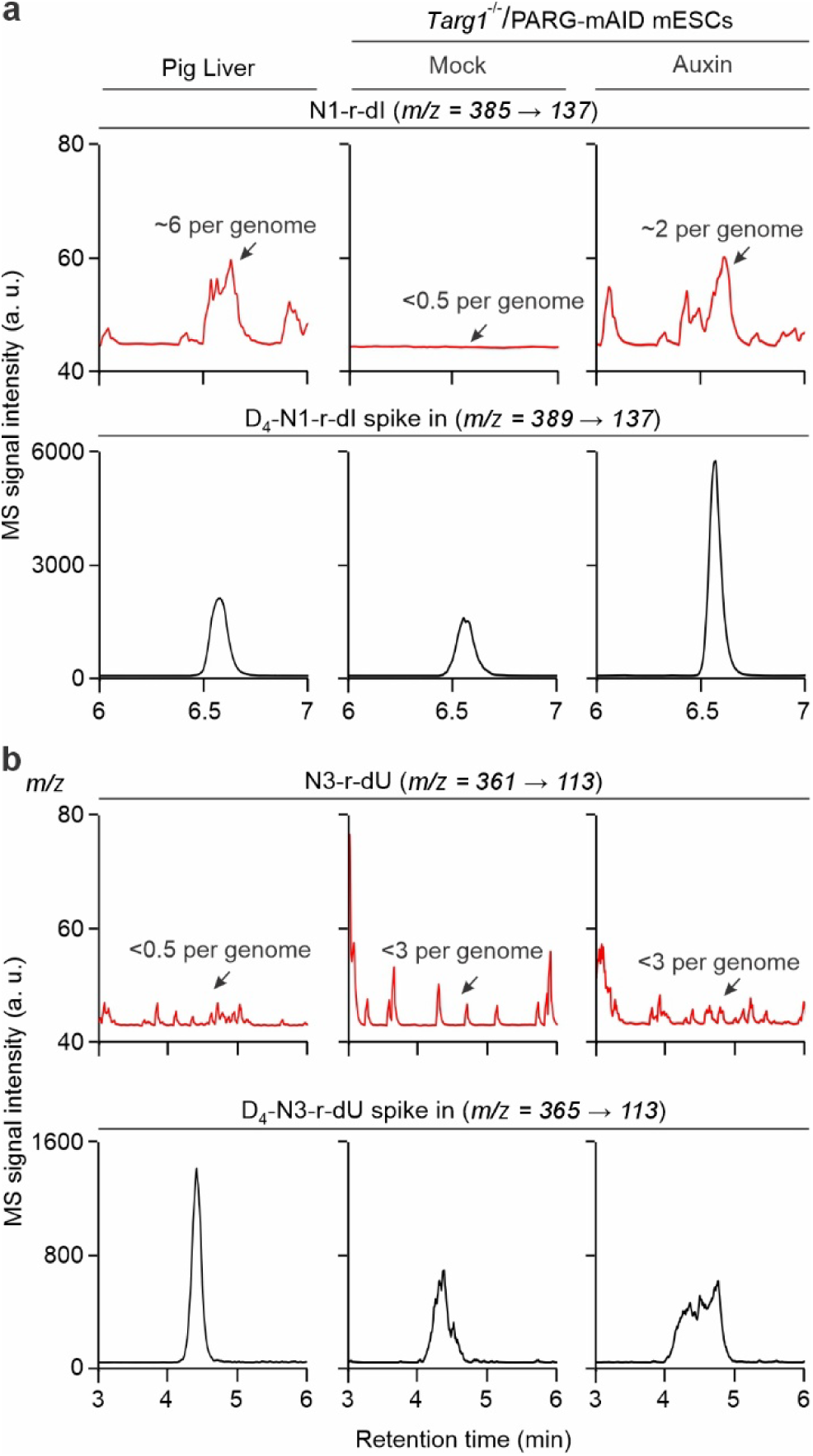
Detection and quantification of DNA base PARylation *in vivo*. **a, b**, LC-MS/MS chromatograms of (a) N1-r-dI (m/z 385 → 137, top) and (b) N3-r-dU (m/z 361 → 113, top) enriched from DNA of pig liver or *Targ1^-/-^/*PARG-mAID mESCs treated or not with auxin for PARG degradation as indicated. Signals for N1-r-dI and N3-r-dU (red lines) are each compared with the LC-MS/MS chromatograms of the respective isotopic standards D_4_-r-dI and D_4_-r-dU (black lines, m/z 389 → 137 and 365 → 113, respectively) spiked-in to the genomic DNA preparations before enrichments. mESCs experiments were done for a single biological replica, pig liver sample was done in triplicates, representative chromatograms are shown.

## Discussion

We report N3-dC PARylation as a novel DNA modification, demonstrate that PARP1 catalyzes the reaction in single-stranded DNA, that N3-PARylated dC is instantly deaminated, and that the PARylation is reversible by TARG1 but not PARG. Moreover, we provide synthetic routes to the relevant diagnostic standards N3-r-dI and N1-r-dU and created inducible PARG, TARG1- mutant mESCs that will be useful for the research community.

Quantifying dC- and dA-PARylation in different oligonucleotides indicates strong sequence-dependence and substrate specificity of PARP1, highlighting the need for future in-depth analyses to identify and characterize optimal substrates. The data reveal selective enzymatic reversibility of DNA PARylation, with PARG confined to PARylated dA. In PARylated proteins, PARG fails to cleave the terminal ADP ribose^15^. We propose that a positively charged nitrogen as in N3-ribosylated dA, yet missing in known ADP-ribosylated amino acids^16^, and in deaminated PARylated bases (**Fig. 1a**) is a prerequisite for PARG to cleave the terminal ADPr. Nevertheless, PARG depletion in mESCs demonstrates dynamic turnover of dA- ADP-ribosylation *in vivo*. Unlike PARG, TARG1 does not require a protonated nitrogen to demodify N-PARylated bases *in vitro*. However, the role of TARG1 for DNA de-modification *in vivo* remains to be determined, as TARG1-deficiency in mESCs did not yield detectable amounts of ADP-ribosylated dA and -dC.

While we corroborate and quantify dA-ADP-ribosylation in genomic DNA, we did not detect dC- ADP-ribosylation *in vivo*. Although *in vitro* PARylation of dA- and dC-residues by PARP1 was equally efficient (**Fig. 2e**), dC- ADP-ribosylation may either be even more transient than dA- ADP-ribosylation and/or restricted to other cell types. Notably, deamination of ADP-ribosylated dC to ADP-ribosylation dU creates a potential source of C→T transition mutations, suggesting that ADP-ribosylated dU is efficiently repaired and therefore fails to accumulate. The situation echoes N6-methyl-dA, a DNA modification that is also maintained at vanishingly low levels in most mammalian cell lines, but with some notable exceptions^17^. Hence, it will be interesting to screen different cell lines and notably cancer cells for DNA ADP-ribosylation . Taken together, we report dC- ADP-ribosylation as a new DNA modification and provide evidence that base ADP-ribosylation is tightly controlled, expanding the repertoire of chemically diverse DNA marks in mammalian genomes. Our findings lay the foundation and provide tools to study the physiological role of DNA base ADP-ribosylation.

## Methods

### Protein expression and purification

#### PARP1

Purification of human full-length PARP1 were performed as described previously^13^.

#### PARG

Codon-optimized full-length human poly (ADP-ribose) glycohydrolase (*PARG*)^18^ was inserted into a Novagen pET-28a(+)-derived vector encoding an N-terminal His_6_-MBP-HRV-3C, and C-terminal Twin-Strep-tag (IBA Lifesciences) fusion protein. His_6_-MBP-HRV-3C-PARG-Twin-Strep-tag was produced in 4 l of the E. coli expression strain BL21-CodonPlus(DE3)-RIL (Stratagene). Protein production was induced with 0.25 mM IPTG overnight at 18 °C. PARG was purified from the cleared cell lysate by a HisTrap HP column (Cytiva), followed by Streptactin affinity chromatography (IBA Lifesciences) according to manufacturer’s instructions including an on-column cleavage of the His-MBP-tag with 3C protease overnight at 4 °C. Purified PARG was stored in 25 mM HEPES-KOH pH 7.4, 500 mM NaCl, 3 mM Desthiobiotin, 1 mM DTT, 5% glycerol.

#### TARG1

Full-length human *TARG1* (*C6orf130*) was inserted into pGEX-6P-3 to encode an N-terminal GST-HRV-3C-fusion protein. Production was essentially as described above but using 0.5 mM IPTG. GST-HRV-3C-TARG1 was eluted from a GSTrap HP column (Cytiva) followed by overnight cleavage at 4 °C with 3C protease, cation exchange chromatography on a HiTrap SP HP column (Cytiva) and size exclusion chromatography on a Superdex 75 16/60 (Cytiva) according to the manufacturer’s instructions. Tagless, purified TARG1 was stored in 25 mM HEPES-KOH, pH 7.4, 150 mM NaCl, 10% glycerol at -80 °C.

### Enzymatic reactions

#### DNA PARylation by PARP1

ADP-ribosylation was performed with 200 nM DNA substrate, 2 mM NAD^+^, 2 µM recombinant PARP1, and 20 µM olaparib, if indicated in ADPR buffer (20 mM HEPES-KOH, pH 7.6, 50 mM KCl, 1 mM DTT, 100 μg/ml BSA) for 60 min at 37 °C followed by phenol/chloroform extraction and ethanol precipitation, using ammonium acetate as a salt. Oligo sequences used for PARylation are listed in **Extended data Table 1**. Control reactions were done with active PARP1 but without DNA substrate, with DNA substrate added after reactions were stopped with addition of phenol/chloroform.

#### DNA dePARylation by PARG and TARG1

PARG and TARG1 treatments were done with 200 nM *in vitro* PARylated DNA substrate and 200 nM PARG or 1 µM TARG1, respectively, in ARG buffer (50 mM Tris-HCl pH 8.0, 2 mM MgCl_2_, 1 mM DTT, 100 mM NaCl) for 2 h at 30 °C. DNA was purified by phenol/chloroform extraction and ethanol precipitation, using ammonium acetate as a salt. Control reactions were done with TARG1 or PARG heat-denatured at 70 °C for 10 min.

### LC-MS/MS for base PARylation

#### Generation of 83mer ssDNA substrates with isotopically labeled nucleotides

Asymmetric PCR was used to generate 83mer ssDNA substrates with heavy isotope labeled nucleotides where in dNTP mixture, one unlabeled dNTP was substituted with the respective heavy isotope-labeled dNTP (^13^C_10_-dGTP, ^13^C_10_-dATP, ^13^C_10_-TTP, ^13^C_9_-dCTP or ^15^N_3_-dCTP, Silantes) as described before^13^. The concentrated ssDNA was used for in vitro ADP-ribosylation and LC-MS/MS analysis as described below. Note, the produced 83mer ssDNA substrates contain a mixture of the respective heavy and natural nucleotide due to usage of unlabeled primer sequences.

#### DNA preparation and LC-MS/MS analysis

Degradation of *in vitro* PARylated DNA was performed as described^13^. An equal volume of isotopic standard mixture ^15^N_3_-dC (Silantes), ^15^N_5_^13^C_10_-dA (Silantes) and synthetic D_4_-r-dI and D_4_-r-dU (see below) was added to the DNA samples and injected for LC-MS/MS analysis. Quantitative analysis was performed on an Agilent 1290 Infinity Binary LC system (Agilent Technologies) using ZORBAX SB-C18 column (Agilent Technologies, 2.1 mm × 50 mm, 1.8 µm) coupled to an Agilent 6490 triple quadrupole mass spectrometer. Elution was performed with 5 mM ammonium acetate pH 6.9 and acetonitrile (ACN), the flow was first linearly increased from 0.3 ml/min to 0.38 ml/min in 0 - 10.5 min, then switched to 0.5 ml/min for 10.5 - 14.5 min, and 0.3 ml/min for 14.5 - 15.5 min. The column was kept at 30 °C. The gradient was: 0 - 3 min, 0% ACN; 3 - 7.5 min, 0 - 5% ACN; 7.5 - 10.5 min, 5 % ACN, 10.5 -12.5 min 5 - 50% ACN; 12.5 - 15.5 min, 0% ACN. The MS source-dependent parameters were as follows: gas temperature 110 °C, gas flow 19 l/min (N_2_), nebulizer 30 psi, sheath gas heater 350 °C, sheath gas flow 11 l/min (N_2_), capillary voltage 2200 V (positive mode), nozzle voltage 0 V, fragmentor voltage 300 V, high pressure RF 80 V and low pressure RF 90 V. Compound dependent parameters are listed in **Extended Data Table 2**. When indicated, DNA was not heat-denatured at 95 °C for 5 min prior to degradation. All quantification were performed using stable-isotope dilution LC-MS/MS. R-dU or R-dI quantification is shown relative to total dA.

### Synthesis of N1-ribosyl-dI (N1-r-dI)

#### 1. Synthesis of compound **3**

A solution of 1,2,3,5-tetra-*O*-acetyl-D-ribofuranose (**Extended Data Fig. 2**, compound **10,** Adamas; 636 mg, 2.00 mmol) in acetonitrile (ACN, 40 mL), was mixed with iron (III) chloride hexahydrate (Adamas FeCl₃·6H₂O; 540 mg, 2.00 mmol). The reaction mixture was heated and kept at 90 °C for 0.5h, with constant stirring. Upon completion, the solvents were removed by SpeedVac to reduce volume. The desired reaction product, 2,3,5-tri-*O*-acetyl-D-ribofuranose (**Extended Data Fig. 2**, compound **3**), was purified by flash column chromatography on silica gel (ethyl acetate(EA)/petroleum ether(PE) gradient, 0-50%EA; Silica-CS(Agela) 40 g cartridge; flow rate: 40 mL/min; with detection**: p-Anisaldehyde/H₂SO₄** TLC stain). After evaporation of extra solvent with SpeedVac, compound **3** was obtained as a syrup (260 mg, 0.942 mmol, 47.1% yield).

Characterization of compound **3**

ESI-MS: *m/z* calculated for C₁₁H₁₆O₈ [M+NH₄]⁺: 294.1; found: 294.4.

¹H NMR (400 MHz, CDCl₃): δ = 5.55–5.07 (m, 3H), 4.42–4.11 (m, 3H), 2.17–2.05 (m, 9H).

#### 2. Synthesis of compound **2**

A solution of 2′-deoxyinosine (**Fig. 2c**, compound **1**; 504 mg, 2.00 mmol) in 2.5 mL of pyridine, was mixed with acetic anhydride (Ac₂O; 1.0 mL, ∼10.6 mmol). The reaction mixture was kept for 16 h at 20 °C, with constant stirring. The excess solvent was removed using SpeedVac. The reaction product 3’,5’-di-*O*-acetyl-2’-deoxyinosine (compound **2**) was purified by flash column chromatography on silica gel (EA/PE gradient, 0–100%EA; Silica-CS (Agela) 40 g; flow rate: 40 mL/min; with UV detection at 254 nm). After evaporation of solvent using SpeedVac, compound **2** (**Fig. 2c**) was obtained as a white solid (457 mg, 1.36 mmol, 68% yield).

Characterization of compound **2**

ESI-MS: *m/z* calculated for C₁₄H₁₆N₄O₆ [M+H]⁺: 337.1; found: 337.3.

¹H NMR (400 MHz, CDCl₃): δ = 12.83 (s, 1H), 8.20 (s, 2H), 6.43 (t, J = 6.4 Hz, 1H), 5.43 (d, J = 5.6 Hz, 1H), 4.41–4.33 (m, 3H), 2.94–2.88 (m, 1H), 2.69–2.65 (m, 1H), 2.14 (s, 3H), 2.10 (s, 3H).

#### 3. Synthesis of compound **4**

3’,5’-di-*O*-acetyl-2′-deoxyinosine (**Fig. 2c**, compound **2**; 336 mg, 1.00 mmol) in 20 mL of toluene was mixed with 2,3,5-tri-*O*-acetyl-D-ribofuranose (**Fig. 2c**, compound **3**; 276 mg, 1.00 mmol), tributylphosphine (n-Bu₃P, Adamas; 607 mg, 3.00 mmol) and 1,1′-(azodicarbonyl)dipiperidine (ADDP, Adamas; 757 mg, 3.00 mmol) on ice-bath under nitrogen atmosphere. The reaction mixture was stirred for 16 h at room temperature under nitrogen atmosphere. The reactants were concentrated with Speedvac, and compound **4** was purified by flash column chromatography on silica gel (EA/PE gradient, 0–100% EA; Silica-CS (Agela) 12 g; flow rate: 20 mL/min; with UV detection at 254 nm). Next, compound **4** was further purified by preparative thin-layer chromatography (pre-TLC) (EA=100%) and subsequent preparative high-performance liquid chromatography (pre-HPLC) ((MeCN/H_2_O (10 mmol/L NH_4_HCO_3_), X-Select 10 µm 19*250 mm, 20 mL/min, UV 254) yielding in the result a product with 70% purity (**Fig. 2c**, Compound **4**). After removing solvent with SpeedVac appearing as solid powder (22 mg, 70% purity, 2.6% yield).

Characterization of Compound **4** ESI-MS: *m/z* calculated for C₂₅H₃₀N₄O₁₃ [M+H]⁺: 595.2; found: 595.0. ¹H NMR (400 MHz, CDCl₃): δ = 8.23 (s, 1H), 7.97 (s, 1H), 6.77 (d, *J* = 3.6 Hz, 1H), 6.41–6.38 (m, 1H), 5.84 (t, *J* = 4.8 Hz, 1H), 5.51 (t, *J* = 5.2 Hz, 1H), 5.44–5.42 (m, 1H), 4.64–4.61 (m, 1H), 4.44–4.31 (m, 4H), 4.22 (dd, *J* = 8.8, 4.0 Hz, 1H), 2.90–2.84 (m, 1H), 2.67–2.63 (m, 1H), 2.16 (s, 3H), 2.14 (s, 3H), 2.10 (s, 3H), 2.05 (s, 3H), 1.95 (s, 3H).

#### 4. Synthesis of compound **5** (N1-r-dI)

A solution of compound **4** (22 mg, 70% purity, ∼25.9 µmol) in 3.0 mL of methanol (MeOH), was mixed with triethylamine (TEA, Greagent; 0.20 mL, ∼1.44 mmol) and deionized water (0.10 mL). The mixture was kept at room temperature, with stirring, for 16 h. After the reaction was finished, the solvent was evaporated with SpeedVac. The crude residue was purified by reversed-phase flash chromatography on a C18 column (Acetonitrile (MeCN)/water gradient, 0–5% MeCN; C18(Agela) 40 g; flow rate: 45 mL/min; with UV detection at 254 nm). After evaporating the solvent with SpeedVac, compound **5** (**Fig. 2c**) was obtained as a white solid powder (1.73 mg, 17.4% yield).

Characterization of the Compound **5**

ESI-MS: *m/z* calculated for C₁₅H₂₀N₄O₈ [M+H]⁺: 385.1; found: 385.4.

¹H NMR (400 MHz, D₂O): δ = 8.46 (s, 1H), 8.30 (s, 1H), 6.52 (d, J = 4.0 Hz, 1H), 6.48 (t, J = 6.4 Hz, 1H), 4.68–4.64 (m, 1H), 4.59 (t, J = 4.0 Hz, 1H), 4.44–4.37 (m, 2H), 4.19–4.16 (m, 1H), 3.99 (dd, J = 12.6, 2.2 Hz, 1H), 3.86–3.76 (m, 3H), 2.89–2.82 (m, 1H), 2.64–2.58 (m, 1H).

### Synthesis of N3-ribosyl-dU (N3-r-dU)

#### 1. Synthesis compound **7**

A solution of 2′-deoxyuridine (**Fig. 2a**. compound **6**, 228 mg, 1.00 mmol) in 1.0 mL of pyridine was mixed with Ac₂O (0.50 mL, 5.29 mmol). The reaction mixture was kept for 16 h at 20 °C, then excess solvent was removed by SpeedVac. The concentrated reactants were purified by flash column chromatography (silica gel; EA/PE gradient, 0–100 % EA; column size: Silica-CS (Agela) 40 g, flow rate: 40 mL·min⁻¹; detection: UV at 254 nm) to give 3′, 5′-di-*O*-acetyl-2′-deoxyuridine (**Fig. 2a**, compound **7**, 206 mg, 0.66 mmol, 66.0 % yield) as a white solid.

Characterization of the Compound **7**

ESI-MS: *m/z* calculated for C₁₃H₁_6_N_2_O₇ [M+H]⁺: 313.1; found: 313.0. ¹H NMR (400 MHz, CDCl₃): δ 8.77 (s, 1H), 7.49 (d, *J* = 8.0 Hz, 1H), 6.30–6.26 (m, 1H), 5.78 (d, *J* = 8.0 Hz, 1H), 5.23–5.21 (m, 1H), 4.39–4.26 (m, 3H), 2.57–2.51 (m, 1H), 2.19–2.10 (m, 7H).

### 2. Synthesis of compound **8**

Compound **7** (62 mg, 0.20 mmol) and 2,3,5-tri-*O*-acetyl-D-ribofuranose (**Extended Data Fig. 1**, compound **3;** 55 mg, 0.20 mmol) dissolved in toluene (5.0 mL) were mixed with n-Bu₃P (121 mg, 0.60 mmol) and ADDP (126 mg, 0.50 mmol) on ice-bath under nitrogen atmosphere. The reaction mixture was incubated at room temperature for 16 h under nitrogen atmosphere, with constant stirring. Then, excess solvent was removed by SpeedVac. The crude reactants were purified by flash column chromatography (silica gel; EA/PE gradient, 0–100 % EA; column size: Silica-CS (Agela) 12 g, flow rate: 20 mL·min⁻¹; detection: UV at 254 nm) to give the desired nucleoside conjugate (**Fig. 2a**. compound **8** with 80 % purity (30 mg, 80 % purity, 21.2 % yield) as a white solid.

Characterization of the Compound **8**

ESI-MS: *m/z* calculated for C₂₄H_30_N₂O₁₄ [M+H]⁺: 571.2; found: 571.0.

#### 3. Synthesis of compound **9** (N3-r-dU)

A solution of compound **8** (30 mg, 80 % purity, approx. 0.042 mmol) in methanol (3.0 mL) was mixed with triethylamine (0.20 mL, 1.43 mmol) and water (0.10 mL, 5.56 mmol). The mixture was stirred at room temperature for 16 h. Then, the excess solvent was removed by speedVac, and reactants were purified by reversed-phase flash chromatography on a C18 column (C18; H₂O/MeCN gradient, 0–5 % MeCN; C-18((Agela), 40 g; flow rate: 45 mL·min⁻¹; detection: UV at 254 nm) to obtain desired N3-r-dU (**Fig. 2a**, compound **9**, 2.87 mg, 18.9 % yield) as a white solid.

ESI-MS: *m/z* calculated for C₁₄H₂_0_N₂O₉ [M+H]⁺: 361.1; found: 361.0. ¹H NMR (400 MHz, D₂O): δ 7.75 (d, *J* = 8.4 Hz, 1H), 6.20–6.15 (m, 2H), 5.83 (d, *J* = 8.4 Hz, 1H), 4.65–4.62 (m, 1H), 4.37–4.33 (m, 2H), 3.96 (dd, *J* = 8.0, 4.0 Hz, 1H), 3.90–3.87 (m, 1H), 3.82–3.63 (m, 4H), 2.34–2.27 (m, 2H).

### Synthesis of D4-N1-ribosyl-dI, and D4-N3-ribosyl-dU

To synthesize D4-N1-ribosyl-dI (D_4_-N1-r-dI), as well as D4-N3-ribosyl-dU (D_4_-N3-r-dU), compound **3** was replaced with its D4 labeled isotopologue, (**Extended Data Fig. 3**, compound **16**)

#### 1. Synthesis of compound 12

D-Ribose (**Extended Data Fig. 3**, compound **11,** 8.0 g, 53.3 mmol, Accela) was dissolved in 80 mL of anhydrous methanol and cooled to 0 °C. Next, Dowex® 50WX8 (Aladdin) was added to this solution, and the mixture was incubated at room temperature, for 24 h with constant stirring. After reaction was finished, the resin was removed with filter by Celite® pad, and the remaining solution was concentrated using SpeedVac. The concentrated reactants were subjected to flash column chromatography (silica gel; methanol(MeOH)/dichloromethane(DCM) gradient, 0–10 % MeOH; column size: Silica-CS (Agela) 80 g, flow rate: 60 mL·min⁻¹; detection: **p-Anisaldehyde/H₂SO₄** TLC stain) to purify methyl D-ribofuranoside (**Extended Data Fig. 3**, compound **12**). After removing the solvents with SpeedVac, the desired compound 11 was obtained as a syrup (6.5 g, 74.3 % yield).

#### 2. Synthesis of compound 13

Compound **12 (Extended Data Fig. 3**, 1.5 g, 9.1 mmol) was mixed with 10 % ruthenium on carbon (Ru/C, Rocknew) (924 mg, 0.91 mmol) and sodium hydroxide (146 mg, 3.65 mmol) in 30 mL of D₂O (Adamas), and kept at 80 °C for 48 h with constant stirring. Then, the mixture was cooled down to room temperature and filtered through a Celite® pad. The filtrate was concentrated with SpeedVac and subjected to flash column chromatography (silica gel; CH₂Cl₂/MeOH gradient, 0–10 % MeOH; column size: Silica-CS (Agela) 25 g, flow rate: 30 mL·min⁻¹; detection: **p-Anisaldehyde/H₂SO₄** TLC stain) to purify methyl D-ribofuranoside-D₄ (**Extended Data Fig. 2**, compound **13**). After removing the solvents with SpeedVac, the desired compound **12** was obtained as a syrup (1.0 g, 65.1 % yield).

#### 3. Synthesis of compound **14**

A solution of compound **13** (**Extended Data Fig. 3**) in 5 mL of pyridine was mixed with 2 mL of acetic anhydride. The reaction mixture was kept at room temperature for 19 h, with constant stirring. Upon completion of the reaction, excess solvents were removed with SpeedVac. The concentrated reactants were purified by flash column chromatography (silica gel; EA/PE gradient, 0–50 % EA; column size: Silica-CS-(Agela) 25 g, flow rate: 30 mL·min⁻¹; detection**: p-Anisaldehyde/H₂SO₄** TLC stain) to give compound **14** (**Extended Data Fig. 3**). After removing the solvents with SpeedVac, the desired compound **14** was obtained as a syrup (1.2 g, 68.6 % yield).

Characterization of the Compound **14**

ESI-MS: *m/z* calculated for C_12_H_14_D_4_O_8_ [M+ NH₄]⁺: 312.2; found: 312.2. ¹H NMR (400 MHz, CDCl₃): δ = 4.90 (s, 1H), 4.29 (s, 1H), 3.38 (s, 3H), 2.11 (s, 3H), 2.10 (s, 3H), 2.06 (s, 3H).

#### 4. Synthesis of compound 15

To a mixture of compound **14** (**Extended Data Fig. 3**, 1.2 g, 4.08 mmol) in 40 mL of acetic acid (AcOH) and 5 mL of acetic anhydride, 0.8 mL of concentrated sulfuric acid was added. The reaction mixture was incubated for 3.5 h at room temperature with constant stirring and quenched by the addition of 400 g of crushed ice. The reactants were extracted with dichloromethane (twice, by 200 mL each). The combined organic layers were washed with H_2_O (200 mL) and saturated aqueous NaHCO₃, (200 mL) then dried over anhydrous Na₂SO₄, filtered, and concentrated. The resulted residue was subjected to flash column chromatography (silica gel; EA/PE, 0–50 % EA; column size: Silica-CS (Agela) 25 g, flow rate: 30 mL·min⁻¹; detection: **p-Anisaldehyde/H₂SO₄** TLC stain) to purify compound **15** (**Extended Data Fig. 3**). After removing the solvents with SpeedVac, the desired compound 14 was obtained as a syrup (655 mg, 49.8 % yield).

#### 5. Synthesis of compound **16**

To a solution of compound **15** (**Extended Data Fig. 3**, 655 mg, 2.03 mmol) in 40 mL of acetonitrile, iron(III) chloride (330 mg, 2.03 mmol) and water (220 mg, 12.2 mmol) were added. The reaction mixture was incubated at 90 °C for 30 min with constant stirring. After cooling to room temperature, the volatile solvents were evaporated with SpeedVac, and the reactants were subjected to flash column chromatography (silica gel; EA/PE gradient, 0–50 % EA ; column size: Silica-CS (Agela) 40 g, flow rate: 40 mL·min⁻¹; detection: **p-Anisaldehyde/H₂SO₄** TLC stain) to purify compound **16** (**Extended Data Fig. 3**). After removing the solvents with SpeedVac, the desired compound **16** was obtained as a syrup (230 mg, 40.4 % yield).

Characterization of the Compound **16**

ESI-MS: *m/z* calculated for C₁₁H₁₂D₄O₈ [M+ NH₄]⁺: 298.1; found: 298.2. ¹H NMR (400 MHz,CDCl₃): δ = 5.54 (s, 0.4H), 5.37 (s, 0.6H), 4.40 (s, 0.4H), 4.28 (s, 0.6H), 2.17–2.05 (m, 9H).

### Cell culture

Mouse E14TG2a embryonic stem cells (mESCs) were obtained from ATCC, number CRL-1821. Cell identity was authenticated by ATCC. E14TG2a cells were cultured on plates coated with 0.1% Gelatin (Millipore) in DMEM supplemented with 15% PANSera ES FBS (PAN Biotech), 2 mM L-Glutamine, 100 µM non-essential amino acids (NEAA, Gibco), 1 mM sodium pyruvate (Gibco), 100 µM 2-mercaptoethanol (Sigma), 1000 U/ml Leukemia inhibitory factor (LIF, Millipore), 100 U/ml PEN-STREP at 37 °C in 5% CO_2_ and 20% O_2_. For PARG depletion PARG-mAID tagged mESCs were treated with 0.5 mM indole-3-acetic acid (auxin, Santa Cruz) for 24h. Mock treatment was performed with 0.5% ethanol (solvent). PARG depletion was confirmed by Western blot analysis using standard procedures and a mouse monoclonal anti-HA antibody (Roche, 12CA5), and mouse monoclonal anti-α-Tubulin antibody (Sigma, T5161) as loading control.

#### Generation of TARG1-deficient and PARG inducible-knockdown (Targ1^-/-^/PARG-mAID) mESCs

To generate mESCs depleted for *Targ1* exon 5 (TARG1 catalytically-deficient cells) 1×10^6^ mESCs were seeded and transfected the next day with either empty or gRNA encoding pX330-U6-Chimeric_BB-CBh-hSpCas9 (Addgene #42230) mixed with the selection plasmid pPGKPuro (Addgene #11349) using Lipofectamine 2000 (Invitrogen) according to the manufacturer’s instructions. Cells were selected with 2 µg/ml puromycin for 6 days, colonies picked, passaged and subjected to genotyping PCR using primers flanking *Targ1* exon 5. Positive clones were expanded for further analysis. PARG inducible knockdown mESCs were generated by C-terminal mAID tagging of PARG essentially as described^19^. Homology donor vectors were constructed with derivatives of plasmids pMK286 (Addgene #72824) and pMK287 (Addgene #72825) in both of which a 3xHA-encoding sequence was inserted 3’ to the mAID-encoding sequence. The osTIR1 expression cassette was inserted into the mouse *TIGRE* acceptor locus as described^20^ using plasmids pX330-EN1201 (Addgene #92144) and pEN396 (Addgene #92142) in which the *PuroR* gene was substituted by *bsr* to enable selection with blasticidin (10 µg/ml final concentration). Single colonies were subjected to genotyping PCR and positive clones were expanded for further analysis. OsTIR1 expression was confirmed by Western blot analysis using a mouse monoclonal anti-V5 antibody (Invitrogen, R960-25, **Extended Data Fig. 5**). Oligonucleotide and homology arm sequences are listed in **Extended Data Table 1**.

### Enrichment of ribosyl-deoxyinosine (N1-r-dI) and ribosyl-deoxyuridine (N3-r-dU) from gDNA

To accumulate endogenous dA- and dC PARylation, *Targ1*^-/-^/PARG-mAID mESCs were auxin or mock-treated as described above to generate two conditions: (i) single TARG1-deficiency, (ii) TARG1 and PARG double deficiency. Genomic DNA was isolated with Blood & Cell Culture Kit (Qiagen) according to the manufacturer’s instructions, with overnight Proteinase K treatment, including an additional RNase A treatment (50 µg RNase A per 300 µg gDNA at 0.3 mg/ml) in 2 mM Tris-HCl pH 7.5, 18 mM ammonium acetate for 30 min at 37°C followed by ethanol precipitation. Degradation of ssDNA was performed with 1400 units of Nuclease S1 per 1 mg of gDNA in 5 mM ammonium acetate pH 5.6, 0.2 mM ZnCl_2_ for 60 min at 37 °C using a total amount of 3.3 and 3.1 mg of DNA from mock or auxin-treated *Targ1*^-/-^/ParG-mAID mESCs, respectively, at 0.7 mg/ml. Then 5 µL of 50 nM D_4_-N3-r-dU, 50 nM D_4_-N1-r-dI standards were added into each sample to compensate for the losses of N3-r-dU and N1-r-dI during the following steps, and to allow for quantifications of gDNA PARylation. Degraded ssDNA was separated from undigested dsDNA by an Amicon ®Ultra-0.5 ml 30 kDa filter unit (Millipore). The ssDNA collected from the flow-through was hydrolyzed with 0.4 U of NP1, 2 U of SVP and 20 U of FastAP per 1 mg of starting gDNA amounts followed by enzyme removal through an Amicon ®Ultra-0.5ml 10 kDa filter unit (Millipore). The flow-through was concentrated ∼20x in a Concentrator plus (Eppendorf) at 4 °C. N1-r-dI and N3-r-dU were enriched on an Agilent 1290 Infinity Binary LC system (Agilent Technologies) using a 4.6 mm × 250 mm,1.8 μm, ZORBAX SB-Aq Rapid Resolution HD (Agilent) by sequential runs with 0.8 mg of starting DNA amount per injection. LC was performed with 5 mM ammonium acetate pH 4.5 and acetonitrile (ACN), the flow was at 0.6 ml/min for 0 - 25 min, then gradually increased to 1 ml/min for 25-40 min, gradually decreased to 0.6 ml/min for 40-44 min and 0.6 ml/min from 44-45 min. The gradient was: 0 - 4 min, 0% ACN; 4 - 25 min, 0 - 20% ACN; 25 - 30 min, 20 % ACN, 30 - 35 min 20 - 0% ACN; 35 - 45 min, 0% ACN at 20 °C. The collection window was from 17.0-17.4 min (N3-r-dU) and 19.9-20.3 min (N1-r-dI), as determined empirically by an analytical run injecting a mix of respective DNA sample with synthetic *D_4_-N3-r-dU and D_4_-N1-r-dI*, which peaks at ∼17.2 min and ∼20.1 min, respectively. The collected fractions were pooled, concentrated to ∼20 µl and analyzed in a single analytical LC-MS/MS run using the conditions described above for analysis of *in vitro* PARylated DNA samples. Compound dependent parameters are listed in **Extended Data Table 2**.

## Acknowledgements

We thank Jonathan K. Kraft for experimental support. This work was supported by the Deutsche Forschungsgemeinschaft (DFG, German Research Foundation) instrument funding – [INST247/768-1 FUGG], and the DFG CRC 1361 (Project 20).

## Author Contributions

Conceptualization, M.U.M., L.S., and C.N.; Methodology, M.U.M., L.S., J.M.S., A.B., M.M.M., S.H., L.F., P.G., G.Y., Q.H..; Formal Analysis, M.U.M., P.G., G.Y., Q.H.; Writing – Original Draft, M.U.M, L.S. and C.N.; Review and Editing, all co-authors; Supervision, C.N.; Funding Acquisition, C.N.

## Competing interest

The authors declare no competing interest.

## Corresponding authors

Correspondence to Christof Niehrs.

## Extended Data Figures

**Extended Data Figure 1:**
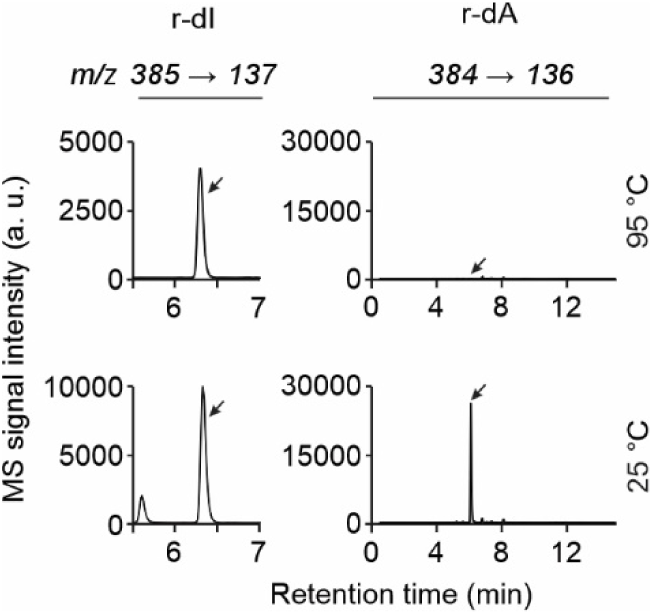
LC-MS/MS chromatograms of reaction products from *in vitro* PARylation. Reaction products with the standard oligo were either processed at 95 °C or 25 °C as indicated before mass spec analysis, and screened for signals with m/z transitions expected for r-dI (m/z 385 → 137, left) and r-dA (m/z 384 → 136, right). Note the prominent signal for r-dA under the non-heated condition (compare to Fig. 1e).

**Extended Data Figure 2:**
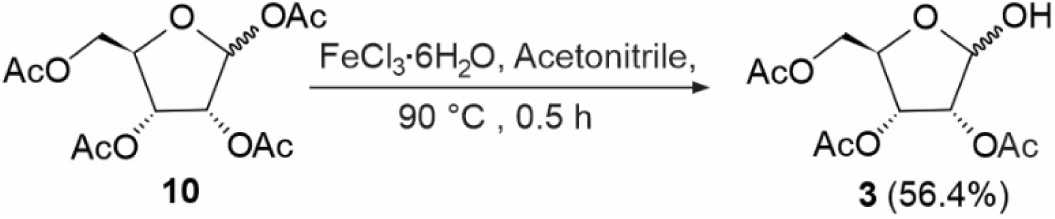
Synthetic route of 2,3,5-tri-*O*-acetyl-D-ribofuranose. Yield is shown in parentheses.

**Extended Data Figure 3:**
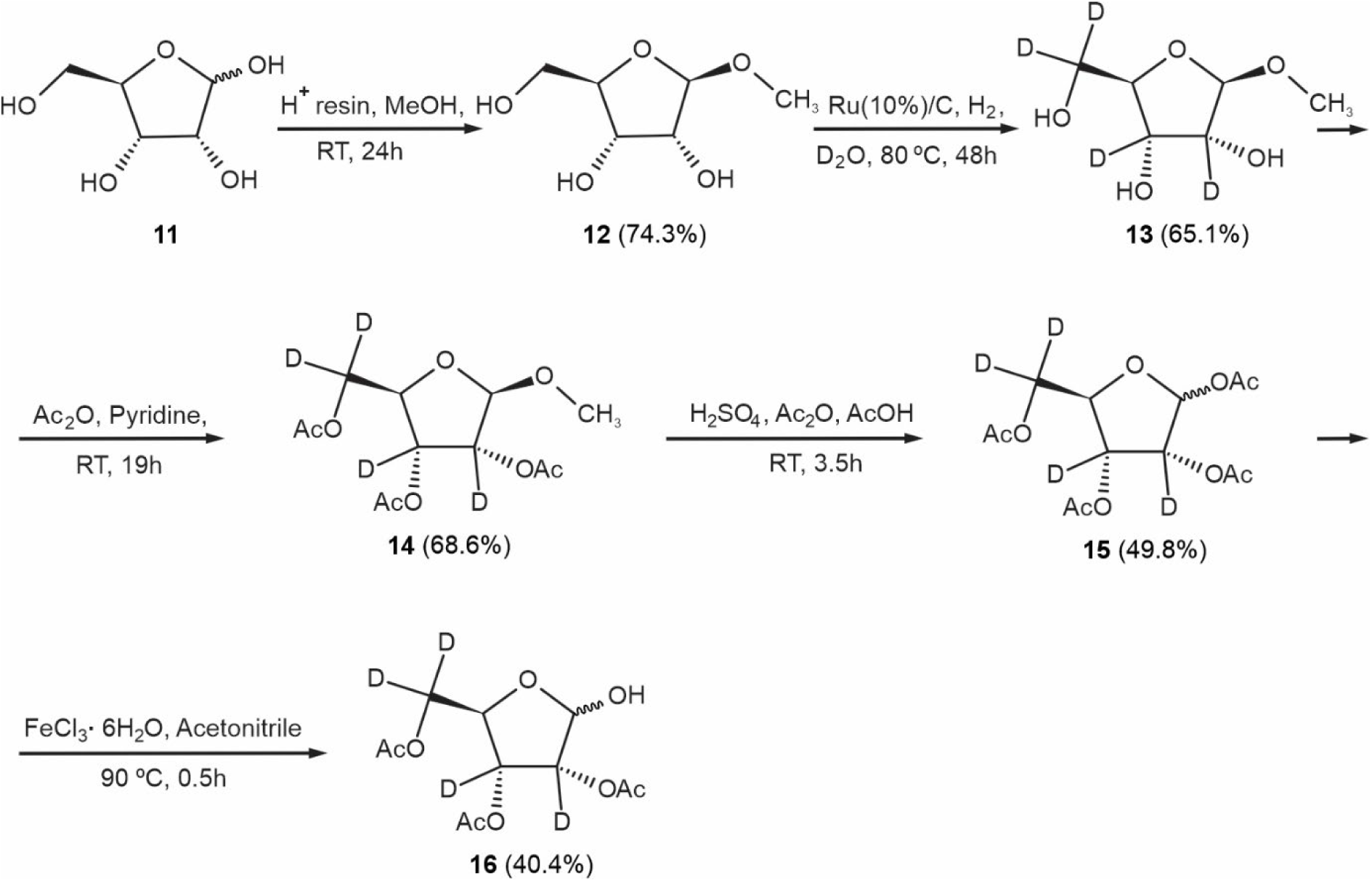
Synthetic route of D4-2,3,5-tri-*O*-acetyl-D-ribofuranose. Yields after each step are shown in parentheses.

**Extended Data Figure 4:**
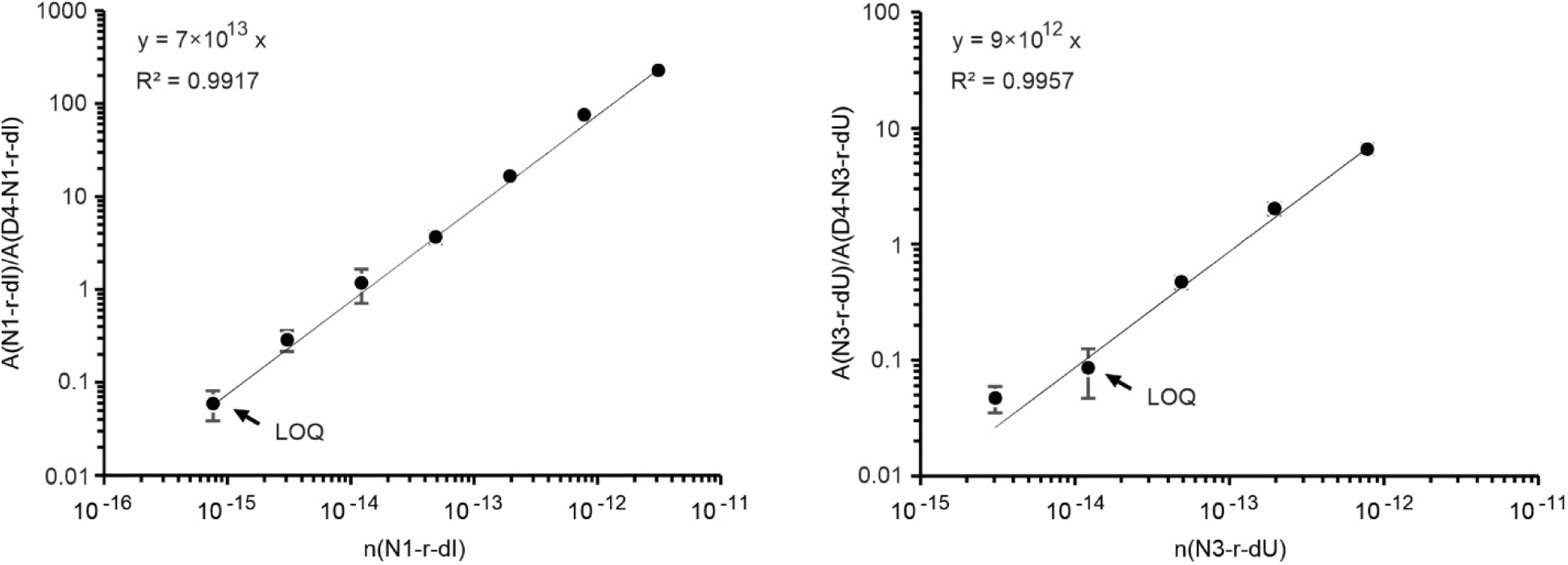
Calibration curves for N1-r-dI and N3-r-dU quantification by LC-MS/MS. The ratio of the signal integrals of either N1-r-dI (left) or N3-r-dU (right) nucleoside standard towards its respective D_4_ isotopic standard is plotted against the amount (n) of nucleoside in number of moles. LOQ, limit of quantification determined as the first point of the linear range. Data are presented as mean, error bars, s.d., n = 3 technical replicates.

**Extended Data Figure 5:**
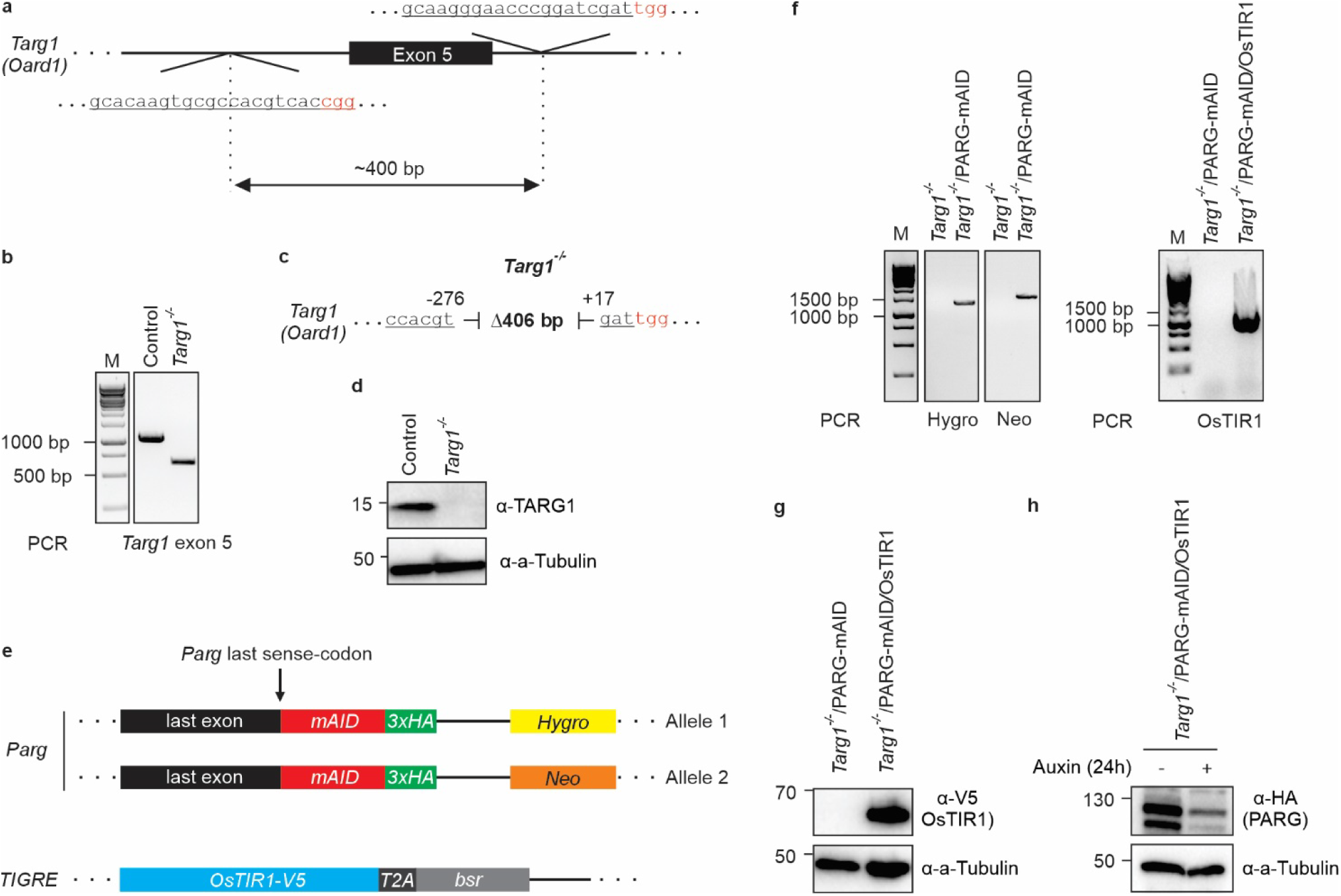
Generation and characterization of Targ1-deficient and PARG inducible-knockdown mESCs. **a**, Scheme of CRISPR/Cas9 targeted *Targ1* exon 5 deletion mutation. To generate genomic deletions, two guide RNAs (target site underlined, PAM motif highlighted in red) were designed to flank a 400 bp region around exon 5 (black bar). Exon 4 encodes the TARG1 catalytic residue K84. **b**, Genotyping PCR of control and *Targ1*-deficient mESC lines. Primers flank a ∼ 1000 bp region around exon 5. Note, the smaller PCR products in *Targ1*-deficient cells indicative of the expected deletion. **c**, DNA sequence analysis of the deleted region around *Targ1* exon 5 in the *Targ1*-deficient mESC line. Size of the deletion is shown within the interrupted sequence with upstream (negative) and downstream (positive) distances in base pairs with respect to the start and end of exon 5. **d**, Western blot for TARG1 in control and *Targ1*-deficient mESC lines. Expected molecular weight for TARG1 is 17 kDa. Alpha (a-) Tubulin served as loading control. Relative molecular weight of marker proteins [x10^-3^] is indicated on the left. **e**, Top, scheme of the *PARG* genomic locus after *mAID-3xHA* insertion 3’-adjacent to the last sense-codon. Note that both *PARG* alleles are tagged with *mAID-3xHA* but with different downstream selection marker genes encoding for either hygromycin (*Hygro*) or neomycin (*Neo*) resistance. Bottom, scheme of the *TIGRE* locus after insertion of *OsTIR1-V5* together with the blasticidin resistance gene *bsr* (separated from OsTIR1-V5 during translation by a T2A peptide). **f**, Left, genotyping PCR of single *Targ1*-deficient and *Targ1*-deficient/PARG-mAID mESC lines. Primers are specific for either the *Hygro* or *Neo* insertion cassette as indicated and flank a ∼ 1300 bp or ∼ 1600 bp region between the respective insertion cassette and the downstream genomic locus. Right, genotyping PCR of *Targ1*-deficient/PARG-mAID mESCs before and after insertion of OsTIR1 into the *TIGRE* locus. Primers flank a ∼ 1200 bp region between the inserted sequence and the downstream genomic locus. **g-h**, Western blots for OsTIR1-V5 in *Targ1*-deficient/PARG-mAID mESCs before and after insertion of OsTIR1 into the TIGRE locus (g), and for PARG-mAID-3xHA in OsTIR1 expressing *Targ1*-deficient/PARG-mAID mESCs treated or not with auxin for 24h (h). Expected molecular weight for OsTIR1-V5 is 67 kDa and for PARG-mAID-3xHA 111 kDa. Alpha (a-) Tubulin served as loading control. Relative molecular weight of marker proteins [x10^-3^] is indicated on the left.

## Extended Data Tables

**Extended Data Table 1:**
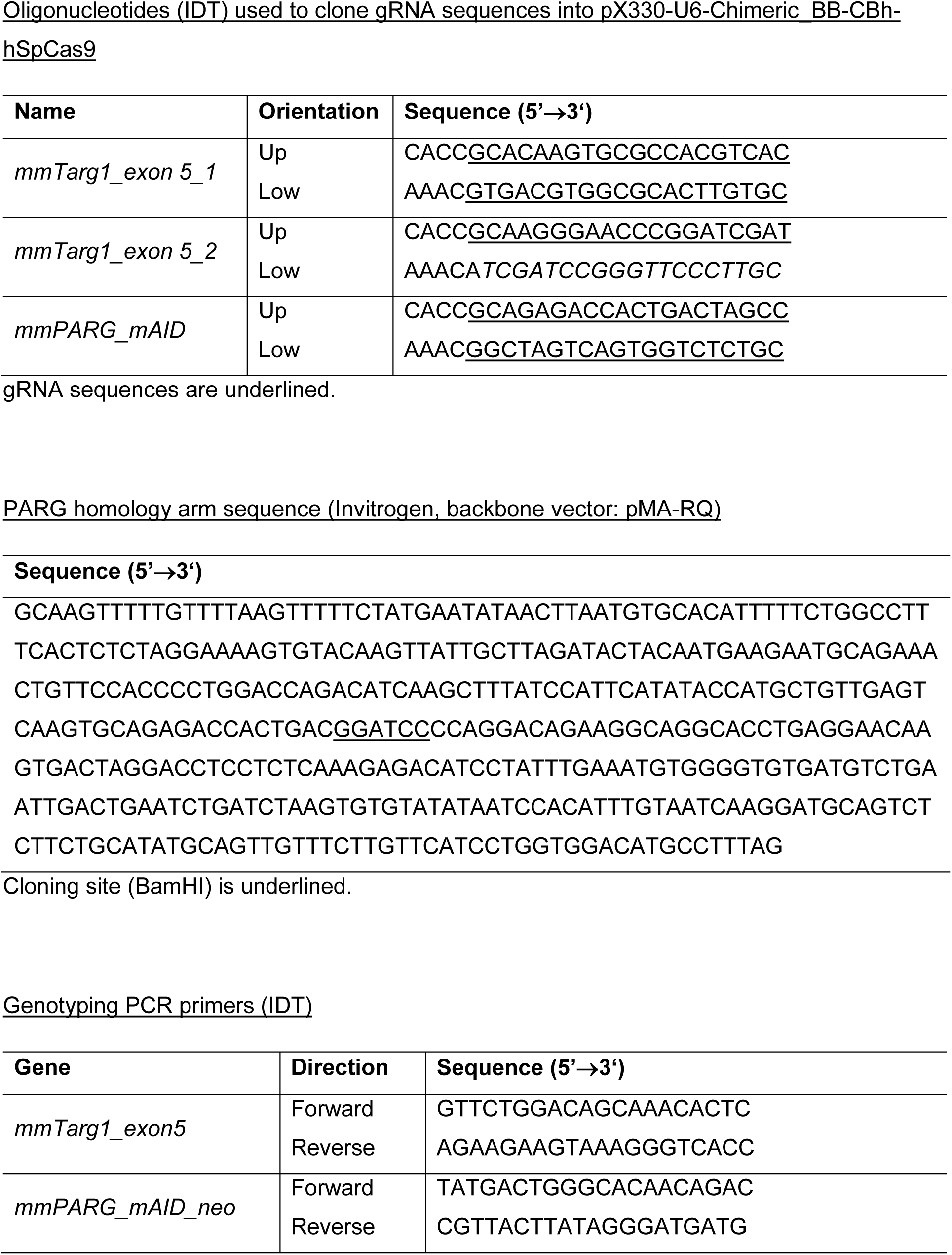
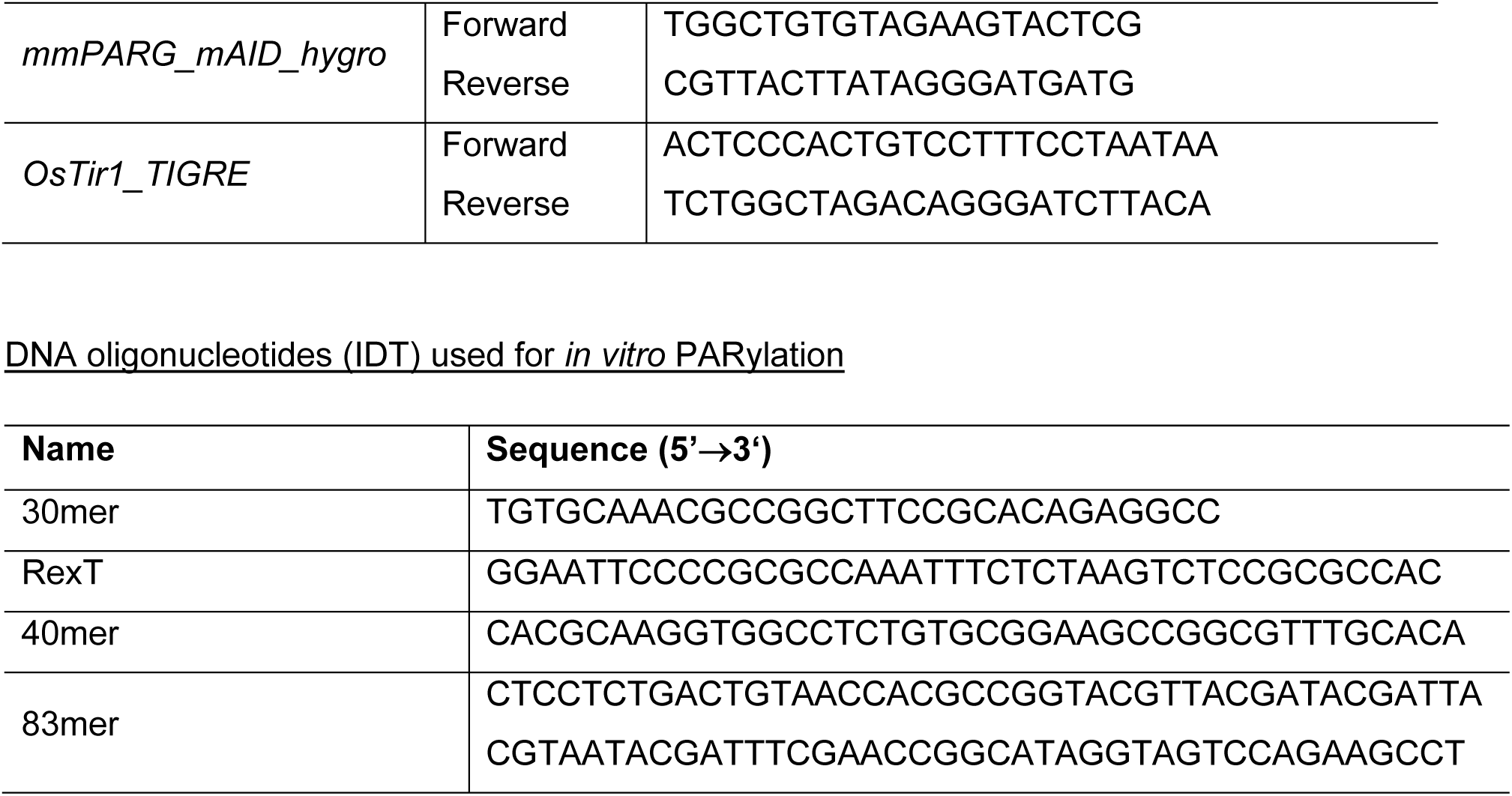
Oligonucleotides used in this study.

**Extended Data Table 2:** MRM transitions used in this study.

| **Nucleoside** | **Precursor ion**<br>**(*m/z*)** | **Fragment ion**<br>**(*m/z*)** | **Collision**<br>**energy** | **Cell**<br>**accelerator**<br>**voltage** |
| --- | --- | --- | --- | --- |
| dA | 252 | 136 | 6 | 8 |
| <sup>15</sup>N<sub>5</sub><sup>13</sup>C<sub>10</sub>-dA | 267 | 146 | 6 | 8 |
| dC | 228 | 112 | 5 | 8 |
| <sup>15</sup>N<sub>3</sub>-dC | 231 | 115 | 5 | 8 |
| N1-r-dI | 385 | 137 | 17 | 5 |
| D<sub>4</sub>-N1-r-dI | 389 | 137 | 17 | 5 |
| r-dU (1) | 361 | 113 | 14 | 6 |
| r-dU (2) | 361 | 245 | 2 | 3 |
| ribosyl-dC | 360 | 112 | 6 | 8 |
| D<sub>4</sub>-r-dU | 365 | 113 | 14 | 6 |
| <sup>15</sup>N<sub>1</sub>-r-dU | 362 | 114 | 14 | 6 |
| <sup>15</sup>N<sub>2</sub>-r-dU | 363 | 115 | 14 | 6 |
| <sup>15</sup>N<sub>3</sub>-r-dU | 364 | 116 | 14 | 6 |
| <sup>13</sup>C<sub>9</sub>-r-dU | 370 | 122 | 14 | 6 |
| <sup>13</sup>C<sub>10</sub>-r-dU | 371 | 123 | 14 | 6 |

MS 1 and 2 resolutions were set to unit for all ions.

## Graphical Abstract

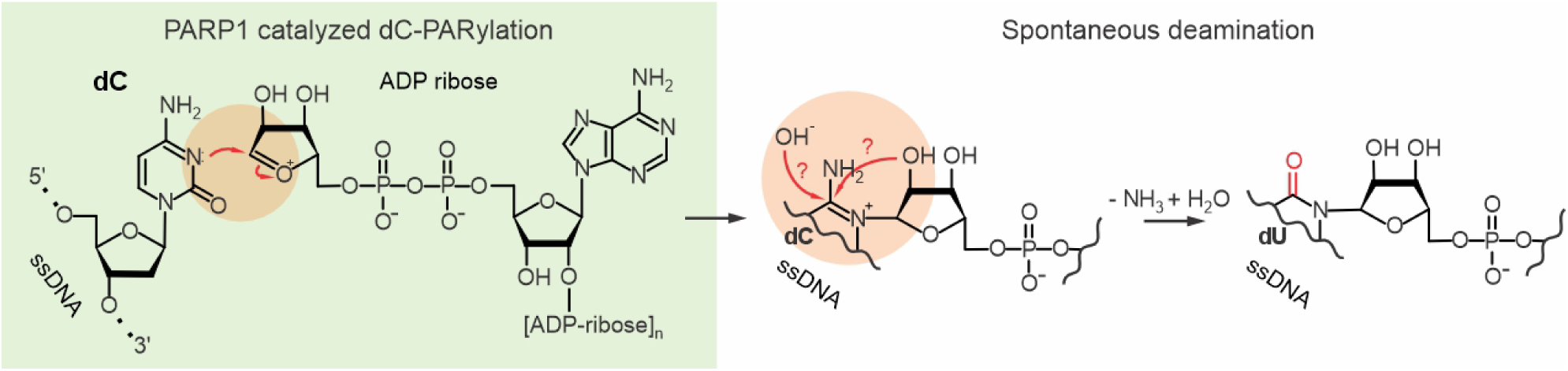

## Notes

### Competing Interest Statement

The authors have declared no competing interest.

